# Estimating Attentional Set-Shifting Dynamics in Varying Contextual Bandits

**DOI:** 10.1101/621300

**Authors:** George Kour, Genela Morris

## Abstract

In this paper, we aim at estimating, on a trial-by-trial basis, the underlying decision-making process of an animal in a complex and changing environment. We propose a method for identifying the set of stochastic policies employed by the agent and estimating the transition dynamics between policies based on its behavior in a multidimensional discrimination task for measuring the properties of attentional set-shifting of the subject (both intra- and extra-dimensional). We propose using the *Non-Homogeneous Hidden Markov Models* (NHMMs) framework, to consider environmental state and rewards for modeling decision-making processes in a varying version of “Contextual Bandits”. We employ the Expectation-Maximization (EM) procedure for estimating the model’s parameters similar to the Baum-Welch algorithm used to train standard HMMs. To measure the model capacity to estimate underlying dynamics, Monte Carlo analysis is employed on synthetically generated data and compared to the performance of classical HMM.

## 1 Introduction

Studies of the neural underpinnings of behavior often assume full knowledge of the represented behavioral parameters. Such knowledge may be achieved by tight experimental control over behavior which allows researchers to correlate neural activity to observable variables that are directly measured or manipulated. However, this approach typically yields artificial, restricted behaviors that are difficult to relate to natural conditions. Conversely, when allowing complex behavior with many degrees of freedom, such as decision making processes involving multiple factors, key elements that contribute to the subjects’ behavior may be latent and therefore the true parameters that are represented by neural activity are unknown and must be inferred from observable data. This is particularly challenging in animal experiments. In such cases, statistical estimation of the parameters underlying the behavior is a necessary first step. In the absence of such statistical tools, researchers must settle on reporting the neural activity and behavioral responses averaged across many trials, neglecting the dynamics of the latent variables. Thus, the aim of trial-by-trial modeling the behavior is twofold: to uncover the “algorithm” by which the animal acts in a certain setup, i.e., to support a candidate theoretical framework of learning and memory underlying the observed behavior; and to compare the neural activity in a certain brain region to the observed behavioral signals in order to learn about its functionality. However, the estimation of these latent variables on a trial-by-trial basis is not trivial and critically depends on the complexity of the hypothesized model and on the specifics of the experimental setup [1]. Therefore, lately, much work was dedicated to estimating internal decision variables by modeling the dynamics of the latent variables and use statistical methods for quantitatively estimating those variables. When the model successfully predicts the subject’s behavior, the values of the model internal variables can be used as a proxy for unobserved decision variables that drive behavior and correlated with neural responses.

Humans and animals are constantly subjected to barrage of sensory information. However, equally considering every aspects of the environment does not allow efficient learning yielding the need to allocate attention to a certain type of stimuli, ignoring details of the others. An attentional set may be considered a cognitive state in which the subject applies the same set of rules for differentiating relevant cues from irrelevant ones in a complex environment [2].

Computationally, an attentional set can be viewed as an efficient state representation formed by the agent while learning a task in a *Markov Decision Process* (MDP) which allows it to behave effectively in the environment. Correspondingly, *Attentional Set-Shifting* (AST) is the process of alternating the attended aspects of the environment to adapt it to new rules. Attentional set-shifting tasks are part of a battery of psychological tests designed to measure attention and cognitive flexibility that has been applied to humans and animals for over half a century. The *Wisconsin Card Sorting Test* (WCST) [3] is a popular test for attentional set shifting in humans. For an artificial agent, such a test would measure its ability to modify the state representation to account for changes in the MDP. Intriguingly, it was found that attentional set-shifting could be impaired in individuals with intact attentional set formation ability. The inability in learning to modify a response when the rules have changed is found in patients suffering from a variety of neuro-degenerative disorders such as schizophrenia, obsessive-compulsive disorders, and Parkinson’s disease [2, 4, 5, 6]. Several types of attentional set-shifting can be measured, *intra-dimensional*, *extra-dimensional*. Intra-dimensional involve shifting attention to a cue that was previously ignored within a dimension (type of cue). Extra-dimensional shifting challenges the flexibility of the attention to shift the attentions o a different type of cue, namely an irrelevant dimension becomes the relevant dimension [2].

We are interested in estimating the attentional set-shifting dynamics of animals in a changing multidimensional discrimination task. This is done by modeling the mechanism underlying a decision-making process using the following experimental setup: A thirsty rat is located at the center of a plus-shaped maze, containing four arms, each enclosed by a door. In each trial, the rat is exposed to sets of two different light and odor cues at two randomly selected doors. To open the door, the rat pokes its nose inside in a hole located at the bottom of the selected door. Once the door is opened, the rat runs into the arm looking for water. The water, if present, is located at the end of the correct arm. Depending on the experimental condition, a specific odor always indicates the correct door (the one with some water), regardless of the light color attached to it, or, in a different condition, a light of a certain color, regardless of its odor. See Figure 1. The discrimination rule can be changed during the experiment to evoke the subject to perform intra and extra-dimensional set shifts. Let us assume that at each trial the rat employs one of the following four policies: odor 1 (O1); odor 2 (O2), light 1 (L1), light 2 (L2). For now, we ignore more complex policies that include more than one independent variable such as *L*1 ˄ *O*2. Since the current policy is not reported by the rat, we aim at estimating the most likely policy employed by the animal in each trial, given the sequence of door selections.

**Figure 1:**
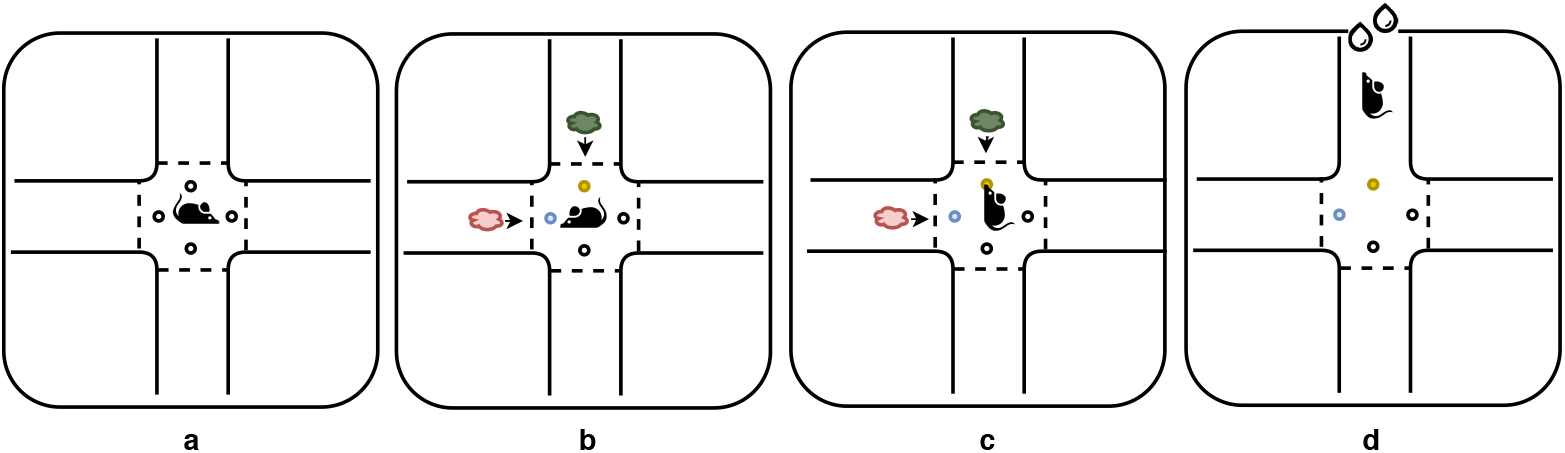
Experimental Setup: Broken lines depict guillotine doors, yellow and blue dots depict LED lights, and green and red arrows indicate the different odors. Selecting the correct door, the water deprived subject receives a reward.

A related behavioral paradigm performed in human subjects is called *Wisconsin cards sorting tests* (WCST). In this task the participant is presented with number of stimulus card and he is asked to match a given card to the other cards according to different criteria. There are several ways to classify the given card (color, shape, number of shapes), and the only feedback is whether the classification is correct or not. The classification rule changes every several trials, and this implies that once the participant has figured out the rule, the participant will start making one or more mistakes when the rule changes. Note while both the WCST and the plus maze are attentional set-shifting tasks, it is easier for the observer in the context of the WCST to identify the current policy of the agent, since in most cases the test card matches only one aspect of each card in the deck. However, in the plus maze, each action of the agent in every state is ambiguous in the sense that it can result from different policies.

In this paper we demonstrate how, under reasonable assumptions, the underlying policies of an agent in a varying contextual bandits can be estimated using NHMMs. The rest of the paper is organized as follows. In Section 2 we present some background that will allow us to formulate and introduce the problem of policy estimation in the context of Reinforcement Learning. We discuss the policy estimation problem in a general sense and also discuss the special case that derived from the nature of the plus maze environment. We show that the policy estimation problem, under some assumptions, is related to HMM. In Section 3 we provide a possible modeling scheme based on an extension of HMM that accounts for changing environment and rewards. In Section 3 we suggest an estimation-maximization algorithm based on a modification of the Baum-Welch algorithm for identifying the proposed model parameters. We illustrate the validity of the proposed model by a Monte Carlo study, in Section 5. Here, we apply our method to simulated data and compare its results with an alternative approach. Related literature is reviewed in Section 6. A summary and future work is discussed in Section 7. Finally, in the appendix, we rigorously derive the modified Baum-Welsh (MBW) algorithm from the expectation maximization process.

## 2 Background

Modeling the dynamics of a system and revealing the its latent states using observable signals emitted by the underlying process, is a fundamental problem in many scientific disciplines. It allows reasoning about the underlying mechanisms that produce the observed signals and prediction of future system states and observations. The *Markov Model* is a key tool in the field of stochastic processes extensively used to analyze the dynamics of time series by extracting meaningful statistics and model the characteristics of the data.

A Markov process is any stochastic process that satisfies the Markovian property, namely, that the future is independent of the past, given the present. Generally speaking, there are four main types of Markov models used in different setups, as shown in Table 1. The differences between the configurations may be schematically viewed as involving two entities: an “environment” and an “agent”. In an autonomous system, in which no agent is involved, the goal of the analysis is to model the dynamics of the environment transitions between the different states; whether the states are fully observable, using *Markov chain* (MC) or obscured by stochastic outputs, using *Hidden Markov Models* (HMM). Modeling HMM requires, in addition to the dynamics of the state transitions, to estimate the dynamics of the observed outputs given the current state. However, in controlled systems, both the agent and the environment are considered. In this case, the mathematical framework aims at modeling a decision-making process in situations where outcomes are partly random and partly under the control of the agent. If it is assumed that the states are fully observable to the agent then *Markov decision process* (MDP) model is considered, otherwise, the modeling is done using *Partially observable Markov decision Process* (POMDP) [7].

**Table 1:** Four types of Markov models.

|  | Fully observable states | Partially observable states |
| --- | --- | --- |
| Autonomous system | Markov chain (MC) | Hidden Markov model (HMM) |
| Controlled system | Markov decision process (MDP) | Partially observable MDP (POMDP) |

Formally, MDP is defined using the tuple 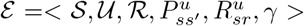, where 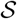 is a finite set of states of size *C*, 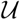 is a finite set of actions of size *M*, 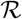 is the set of possible rewards. In MDP, the states are assumed to be Markovian, namely, that the environment’s response at time *t*+1, depends only on the state and action at time *t*. 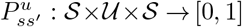 is the state transition probability distribution, 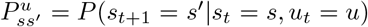, where *s*, 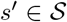 and 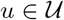. 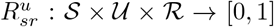 represents the a probabilistic reward function, 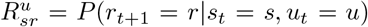, and *γ ∈* [0, 1] is the reward temporal discount factor.

The main problem of MDP (and POMDP) is to find a policy function 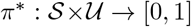 for the agent to use in order to maximize the accumulative reward, 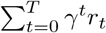. If the state transition probabilities and the reward expectation function are given and the number of states is not extremely large, linear programming or dynamic programming methods can be used to find the optimal policy. However, if these functions are unknown to the agent, the agent must explore the environment and receive sufficient knowledge about the environment to find a good policy using *Reinforcement learning* (RL) techniques [8].

It is important to note that the plus maze environment described above is an instance of the *contextual bandits* problem [9], a special case of the general MDP described here. Contextual bandits can be viewed as a one-step RL, in which the next state of the environment is independent of the previous state and of the agent’s action. In *varying contextual bandits* the underlying distribution of reward remains unchanged over (possibly short) epochs and shifts at unknown time instants.

## 3 Context Aware Hidden Markov Model

Given an agent behaving in a varying contextual bandits environment; we are interested in modeling the dynamics underlying it’s decision process. For instance, imagine an interaction between an agent and the environment where in each game the environment presents two options for the agent. Each option is marked with a number of exclusive and exhaustive cues from several categories, taken from a randomly selected permutation of cues. The agent, whose goal is to maximize its reward, selects one of the two options in each game. Given a sequence of options and selections, can an observer estimate the strategy followed by the agent in each game? For the rest of this paper, we will refer to the agent whose behavior is being inferred as the *agent*, and the entity who seeks to infer a model for these decisions as the *observer*.

In the general sense, one can think of this setting in RL terms as the ***policy estimation problem***, in which the goal is not to find the best policy to accumulate reward but rather to estimate the policy currently used by the agent based on its observed behavior. Formally, given a trajectory of states, actions and rewards, denoted by 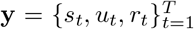; and a class of possible policies that can be employed by the agent 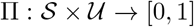, the state transition and reward probabilities of the MDP ε, we are interested in finding the most likely sequence of policies 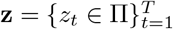 used by the agent during its interaction with the environment. In this paper this dynamics is viewed as two parallel interacting probabilistic processes, one controlling the environment behavior and the other controlling the agent behavior.

However, note that in the general case, the agent’s policy changes continuously, constantly refined in response to feedback. The policy update procedure can be performed in countless ways. The update can be performed after each step of interaction or, in an episodic setup, after one or more episodes is completed. Therefore, with no additional restrictions and assumptions on the set of policies the agent considers and on the policy update technique, the problem of identifying *z*_*t*_ is difficult.

We will impose several restrictions to make the problem statistically identifiable and computationally feasible. We shall require some notation for sequences over finite alphabets. *x*_*1:n*_ will denote a string of length *n* where *x*_*i*_ will denote the the *i*’th element of a sequence, and *x*_*i:j*_ will denote elements *i* to *j*, inclusive where the indices will often be time indices. In addition, use bold fonts to represent vectors and capital letters to denote matrices.

### 1. Finite state observable MDP

We assume that the observer is aware of all the environmental features considered by the agent and that the set of environmental states is discrete, 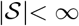. Specifically, let as assume that 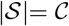

### 2. Discrete set of strategy

While in the general case, a policy is an arbitrary probabilistic mapping between state and action, we assume that the agent employs a fixed number of policies, which we call *strategies*; 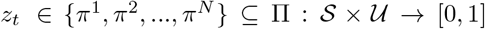. In the general case, the strategies and the transition dynamics between them are unobserved and the goal of the model is to identify these strategies and estimate the transition between them. Note that in specific cases, the observer may have access to or may assume the set of policies employed by the agent. In such cases, the goal of the model is to find the transition dynamics between the different strategies.

### 3. Reward dependent Markovian policy transition

We assume that the policy transition is a first order Markov process that depends only on the previous policy and current reward. Formally:

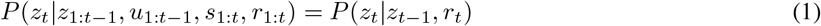

While this is may not be true in the general case, as policy updates may depend not only on the last policy and current but also on previously employed policies, we claim that this assumption is not too “harsh” due to the fact that it is true for state-of-the-art techniques to train artificial agents. This is because, in these approaches, the update step is performed relative to the last policy employed discarding previous policies. Additionally, the assumption that the policy update step is affected by the last policy employed and last received reward fits the *“law of effect”* principle [10], which states that if an action was followed by negative reward (or no reward), the probability that the same action will be repeated in the same state is reduced. Conversely, if an action yielded a positive reward in a given state, the probability that the same action will be repeated will increase.

### 4. Varying Contextual Bandits

The contextual bandits can be considered as a one-step RL (the next environment state does not depend on the previous state nor on the agent’s previous action). therefore we can assume that

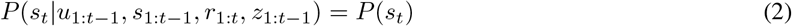

In addition, while we restrict the reward function to return a discrete reward, namely, 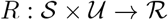 where 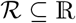 we allow probabilistic varying reward, namely that the reward function of the environment may be a probabilistic function that may change during the interaction.

Formally, the dynamic process we call *Context-Aware Hidden Markov Model* (CA-HMM) can be described by the tuple:

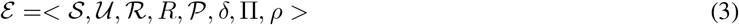

where the first five components 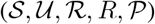 are related to the environment and he last three (*δ*, Π*, ρ*) are related to the agent’s behavior. 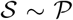 is a finite set of environment states distributed according to distribution 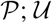 is a finite set of actions, 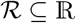 a fixed set of possible rewards, and *R* is the reward function, 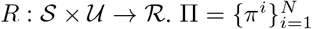 is a discrete set of policies, *π^i^*: *S × U →* [0, 1]. *ρ* is the probability of each *π^i^ ∈* Π to be employed by the agent in the first trial, and 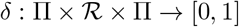 is the transition function between policies following the reward.

Following those assumptions the temporal dependency between the observed and unobserved variables (which is refereed to as the *Markov law of motion* in econometric literature [11]) factors into:

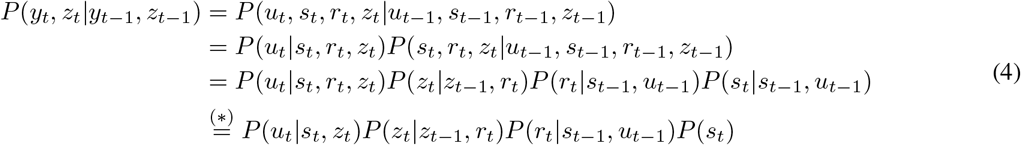

Transition (*) is a result of incorporating our modeling assumptions. The first two terms in the last expression depends on the agent dynamics and are referred to as the conditional actions probability and the the conditional transition, respectively. The remaining terms are dependent on the environment dynamics will be referred as the environment law of motion. This resulting dependency diagram is visualized in Figure 2. Denoting by *θ* the parameters of the model governing the agent behavior, the complete-data likelihood *P* (**y**, **z**; *ε, θ*) can be factorized as follows:

**Figure 2:**
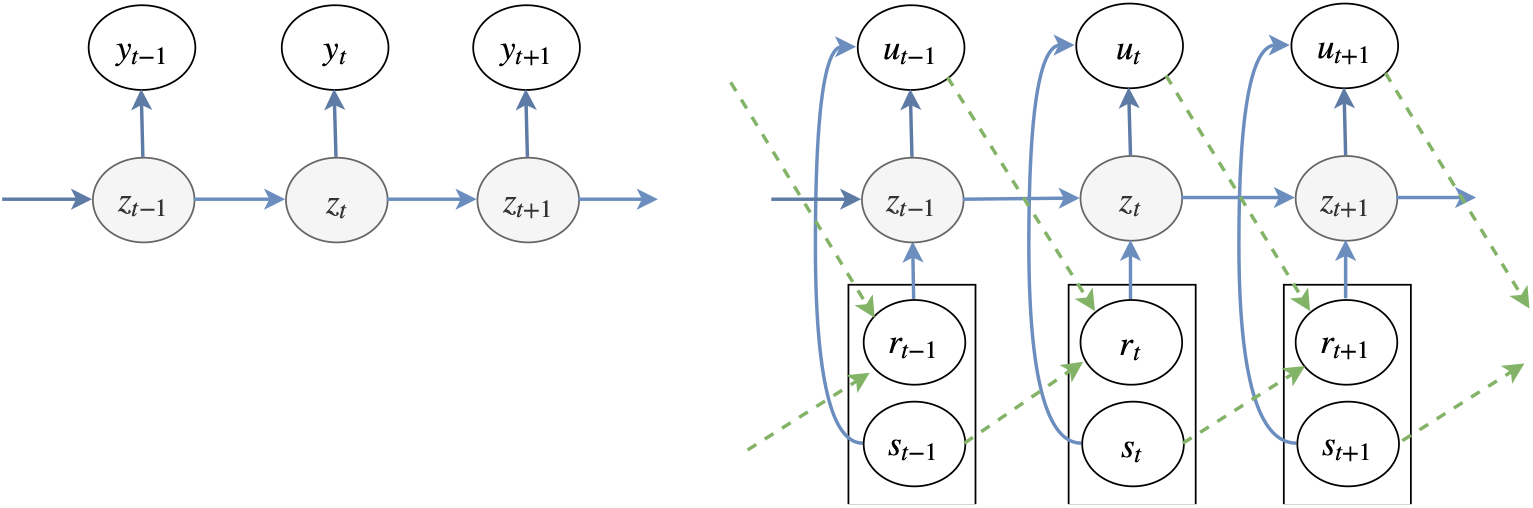
A comparison between the dependency diagram of the standard HMM (Left) and the CA-HMM model (Right). Shaded circles represent unobserved variables. Blue arrows represent probabilistic dependency between variables governed by the agent dynamics (*θ*), and the green arrows represent dependency controlled by the environment dynamics (ε). Continuous lines are unknown (to the observer) dependencies to be estimated by the algorithm. Dashed lines are dependencies known to the observer but not to the agent, thus need not to be estimated by the observer.

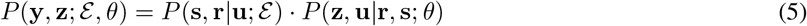

#### Proof

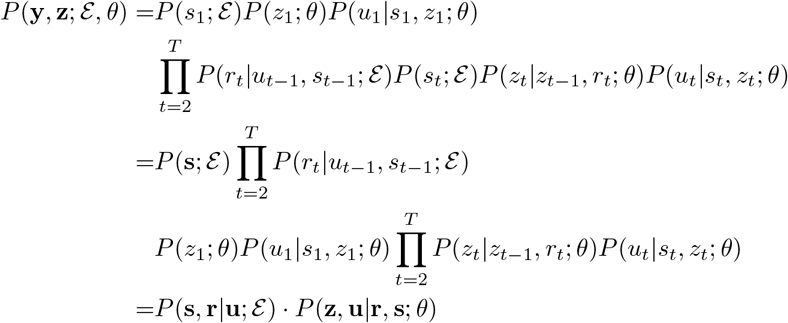

First, note that the complete data likelihood factorized to two terms that the first depends on the environment parameters *ε*, and the other depends on the agent parameters *θ*. Second, note that the closely related to the complete data likelihood of HMM where the sequence of policies corresponds to the sequence of the latent states in an HMM, and the trajectory corresponds the observed emissions’ sequence. However, there are two main differences between estimating policies (under the assumptions we imposed) and HMM: (1) Transitions are reward-dependent transition and (2) Emissions depend on the state of the environment.

## 4 Parameters Estimation: Modified Baum-Welch algorithm

In the following we propose a method for estimating the sequence of policies employed by the agent given the trajectory of the observed variables. The model parameters (*θ*) to be identified include: (1) the probabilities controlling the transition between policies (*δ*); (2) the parameters of the *N* different policies (i.e. the set Π); and (3) the probability of employing each policy on the first trial (i.e., *ρ*).

First, let us specify the parameters related to the transition between policies. We denote by 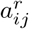 the probability of transition from policy *π^i^* to policy *π^j^* after receiving reward *r*, formally 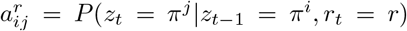. Second, we specify the parameters controlling each of the *N* different policies employed by the agent.

Let us re-arrange the model’s parameters in variable matrices compatible with the transition and output matrices of HMM. Suppose that *A^r^* is a matrix specifying the transition probabilities between policies in the presence of reward *r*, namely 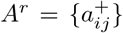, where 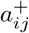 denotes the probability of transition between policy *i* to policy *j* given reward *r*. Without loss of generality let us assume two possible reward values: 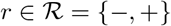 where ‘+’ denotes the presence of a positive reward and ‘*−*’ otherwise.

Recall that the observed information in our model are the actions of the agent and the environment state at each step *t*. Since the agent’s actions depend not only on the current policy but also on the current environment state, we use separate output matrix for each state. Therefore, we define 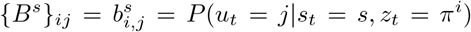, namely the cell *i, j* in the matrix *B^s^* denotes the probability of policy *π^i^* performing action *j* in state *s*. Thus, *B*^1^*, B*^2^*, …, B^c^* denote the output probabilities of the different policies in each state of the environment. Note that the row *i* of matrix *B^s^* denotes the distribution of actions in policy *i* in state *s*.

Following the result of Equation 3, the data likelihood with respect to the parameters *θ* and ε can be written as:

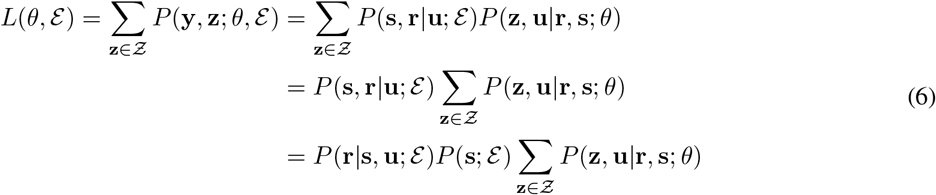

where 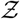 is the set of all possible trajectories of policies of length *T*. However, the term *P* (**r|s**, **u**; ε)*P* (**s**; ε) does not depend on *θ*, therefore, given the observed variables **s** and **r** the likelihood with respect to *θ*, can be written as:

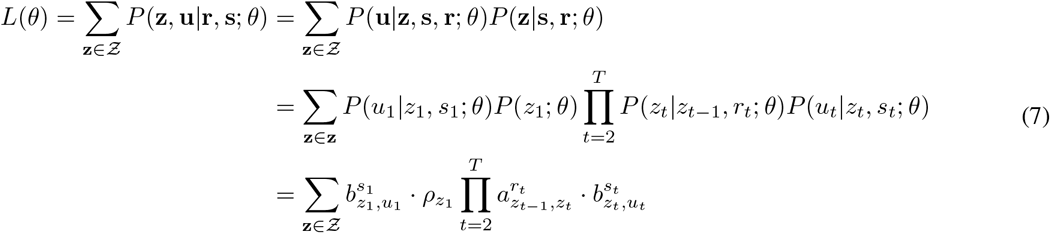

The Baum-Welch (BW) algorithm [12, 13] is a well established Expectation-Maximization method for estimating the parameters of HMM that was developed in the late 1960’s [14]. We modified the BW algorithm to estimate the parameters of the CA-HMM by adapting both the forward-backward and the update procedures to account for environmental states and for rewards.

We will represent the set of the trials in which the animal received a reward as Ψ^+^ and the complementary set of trials as Ψ^*−*^. Formally Ψ^+^ = {*t|w_t_*= +} and Ψ^*−*^ = {*t|w_t_* = −}. Let us denote Ω^*c*^ = {*t|s_t_* = *c*} as the set of trials in which the state is *c*.

### Forward Procedure

Let *α_i_*(*t*) = *P* (*u*_*1:t*_, *z*_*t*_ = *π*^*i*^|*r*_1:*t*_, *s*_1:*t*_; *θ*) the probability of observing actions *u*_1_*, …, u_t_* and being in policy *i* at trial *t*. *α_i_*(*t*) can be computed recursively as follows:

- 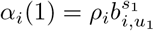
- 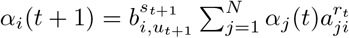

### Backward Procedure

Let *β_i_*(*t*) = *P* (*u*_*t*+1:*T*_|*z*_*t*_= *π^i^*, **r**, **s**; *θ*) the probability of the observed partial action sequence starting at time *t* + 1 given that the policy at time *t* is *π^i^*. We calculate *β_i_*(*t*) as:

- *β*_*i*_(*T*) = 1
- 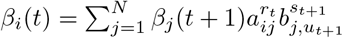

For each trial *t* and every state *i* we first calculate *γ_i_*(*t*) which is the probability of being in state *i* at time *t*, given the observed sequence **y** and the parameters *θ*.

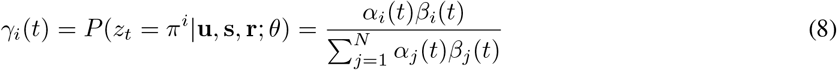

Then we calculate *ξ*_*ij*_(*t*), which is the probability of employing policy *π^i^* and *π^j^*at times *t* and *t*+ 1, respectively.

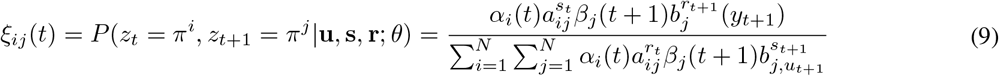

### Update Procedure

Given *γ_i_*(*t*) and *ξ_ij_*(*t*) we update the transition and output matrices. The transition matrix, reflecting the expected fraction of transitions from policy *π^i^* to policy *π^j^* of all transitions from policy *π^i^* is updated as follows:

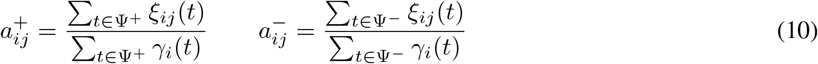

Let *I*(*x*) indicate the indicator function which is 1 if *x* is true and 0 otherwise. The observation probabilities are calculated independently for each environment type, as follows:

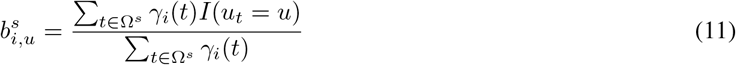

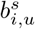 is the estimated probability of taking action *u* when employing policy *i* and state *s*.

The initial state probabilities can be calculated for each state using the parameter *γ_i_*(1).

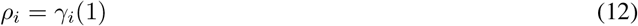

which is the expected frequency spent in state *i* at time 1.

## 5 Model Validation

To empirically test the validity of the inference algorithm, we use synthetic data that contains four underlying policies, two states and two possible actions at each state, with a reward given to a one action in every state, while the other state is never associated with a reward. To measure the precision of the policies’ estimation for different sequence lengths, we generated sequences with lengths ranging between 10 and 1000. We are particularly interested in testing the algorithm performance on parameter complexity and values close to the plus-shaped maze. Therefore, the parameters of the synthetic data generator was set accordingly. In the following, we use the terminology of the plus-shaped maze described in the previous sections. In this case, ignoring the door direction, each trial is one of two different states: one is *s*_1_ = (*O*_1_*L*_1_*, O*_2_*L*_2_), namely that the first door is associated with the cues *O*1 and *L*1, and the second with the complementary cues. The second state is (*O*_1_*L*_2_*, O*_2_*L*_1_) which represents the state in which the first door is associated with *O*_1_ and *L*_2_ and the second with the other cues.

The parameters of the generative process are as follows:

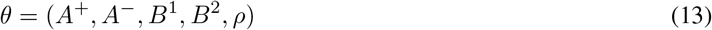

where

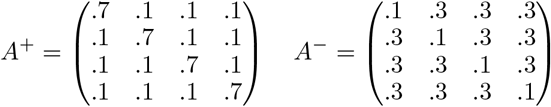

and

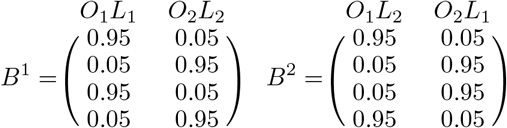

Assuming *L*_1_ is the correct cue, that is always rewarded and *P*(*s*) = 0.5*∀s ∈ S*.

In Figure 3 we compare the convergence pattern of the MBW algorithm assuming CA-HMM dynamics to the convergence of the Baum-Welch algorithm assuming HMM dynamics. We use the following parameters for the HMM: *θ_HMM_* = (*A*^+^*, B*^1^*, ρ*). Each row in the transition matrix is a distribution, thus, to calculate the differences between the true and estimated matrices, we aggregated the total Kullback-Leibler (KL) divergence [15] values of the matrices rows and averaged over the two transition matrices, rewarded and unrewarded. When considering the different policy matrices, we similarly averaged over the matrices of the different states. In Figure 3 we can see that both models converge to the same KL distance between the estimated and real transition probabilities, and that the convergence rate of the MBW algorithm in the CA-HMM model is comparable to that of the Baum-Welch algorithm the HMM model. Note that for the given parameters, one can obtain a good estimation (on average) of sequences of length 300. In our experimental setup, rats perform approximately 100 daily trials, and each rat is trained for 5 − 8 days. Assuming that the policies and the transition probabilities between consecutive days are stationary, the amount data extracted from our experiments should be sufficient to achieve a reasonable parameter estimation.

**Figure 3:**
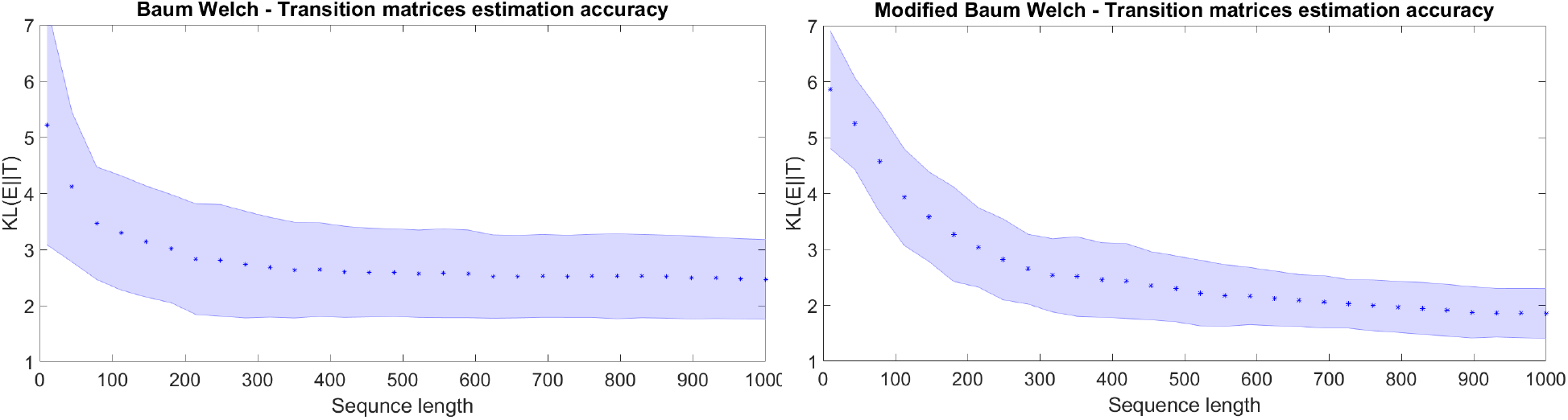
Comparing the accuracy convergence of the estimated transition matrices using BW for HMM and MBW for CA-HMM. The stars represent the mean value of 50 runs and the blue area represents the standard deviation.

In order to show that the MBW algorithm outperforms the BW algorithm in estimating the parameters in the CA-HMM setup, we compared the estimation performance of MBW to BW on data generated using an CA-HMM data generator. The testing methodology is described in Figure 4. In the first stage of the flow, we randomly generate training and test data, each containing sequences of states, actions and rewards, based on the parameters specified in Equation **??**. The training sequence is used by both algorithms to estimate the corresponding model parameters. Following the training stage, the estimated parameters are used to find the underlying policy sequence of the testing sequences. Finally, we compare the policy hit rates on a trial-by-trial basis of both BW and MBW. A modified version of the Viterbi algorithm [16] was used when CA-HMM is considered, to account for the observed rewards and state configurations at each time step. In Figure 5a we show the mean hit-rate result for each sequence length calculated over 50 repetitions. Note that the test sequence length is always 200. We set the tolerance parameter to 0.01 for both BW and MBW. This parameter specifies the convergence indication of the iterative estimation process. The guess transition and emission matrices were generated based on the true probability obfuscated using 50% Gaussian noise. To make the results of the BW and the MBW comparable, the initial guess parameters for the BW algorithm are set as the? average of the rewarded and the unrewarded transition matrices. The output matrices were handled similarly. We can see that the MBW substantially outperforms the BW algorithm in estimating the hidden state given the observed data. Using the same paradigm, Figure 5b shows the posterior probability of a fixed probe sequence as a function of the length of the training sequence.

**Figure 4:**
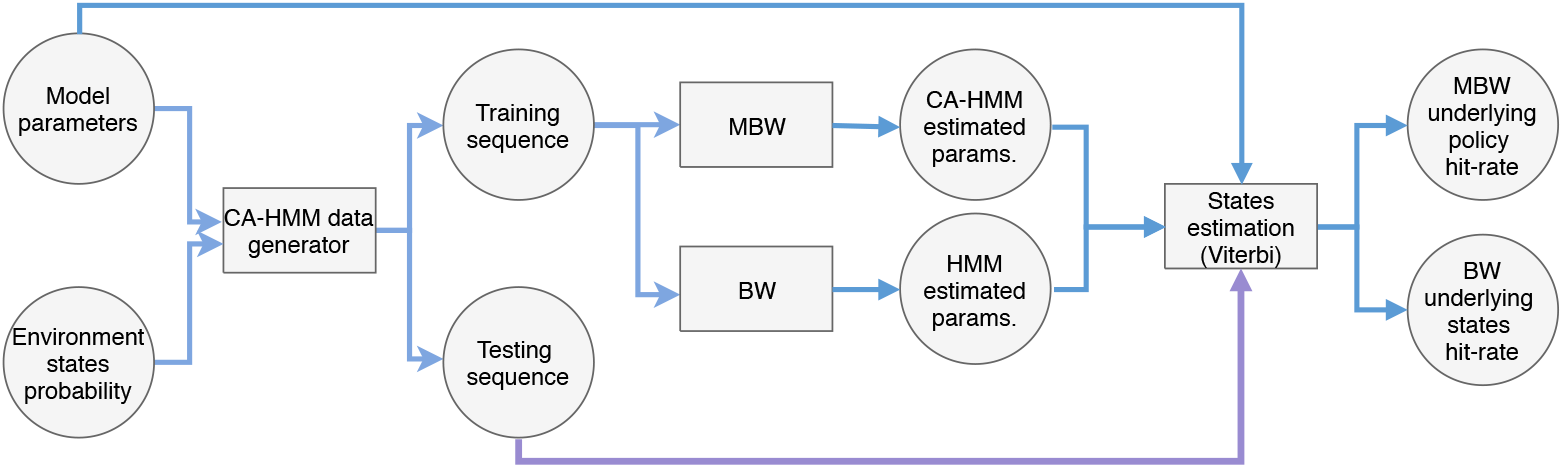
Experimental Flow (single iteration).

**Figure 5:**
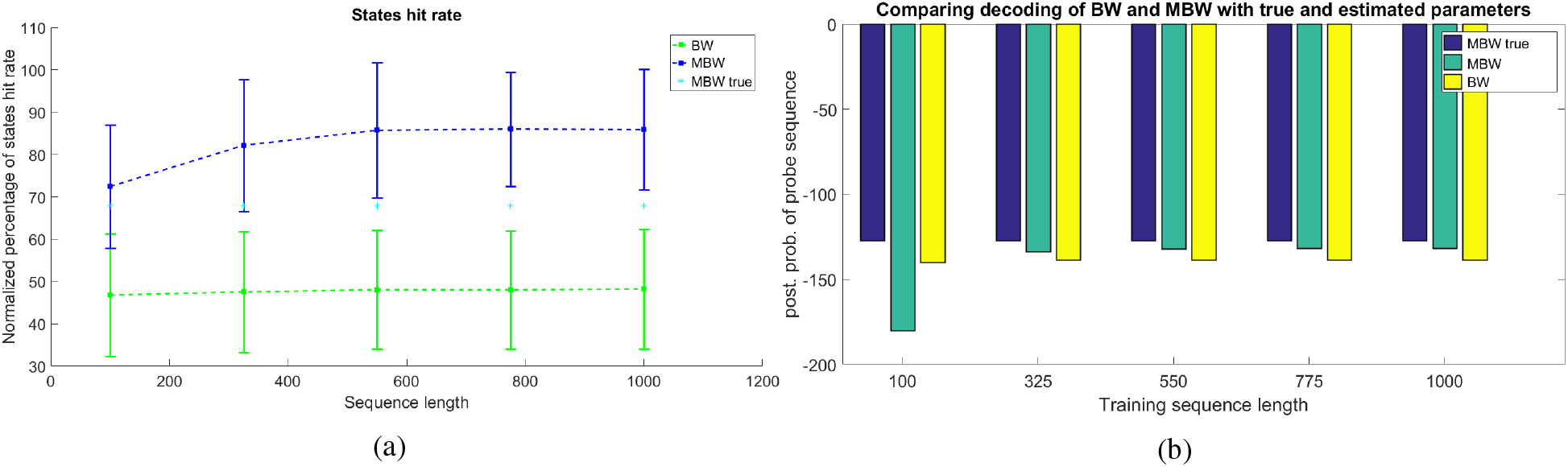
(a) Comparing the state hit rate of MBW and BW using the corresponding Viterbi algorithm. The plot shows the normalized percentage of correct state (policy) sequence estimation versus the length of the learning sequence. The probe sequences contain 200 artificially generated trials. The hit rate of the real parameters are shown (asterisk) to illustrate the maximum possible hit rate. (b) Posterior log probability of a single probe sequence (containing 200 trials) using a model trained by BW and MBW. The blue bar is the log probability of the probe sequence using the true model parameters.

In the following experiments, we attempt to provide evidence for the stability of our model by showing that (1) the estimation precision increases with the accuracy of the initial guess?; and (2) the number of iterations needed for convergence decreases with the accuracy of the initial guess.

In Figure 6 we show the discrepancy of the estimated and true transition matrices, versus the size of the noise factor which represents the amount of noise entered to the perfect guess. The noise was introduced only to the transition parameters - the guess of the policies is perfect. We can see that the algorithm estimation error and the number of iterations needed are almost linear with the amount of noise. In this experiment there were 50 repetitions for each noise factor, the tolerance parameter is 0.01 and length of the training sequence is 1000.

**Figure 6:**
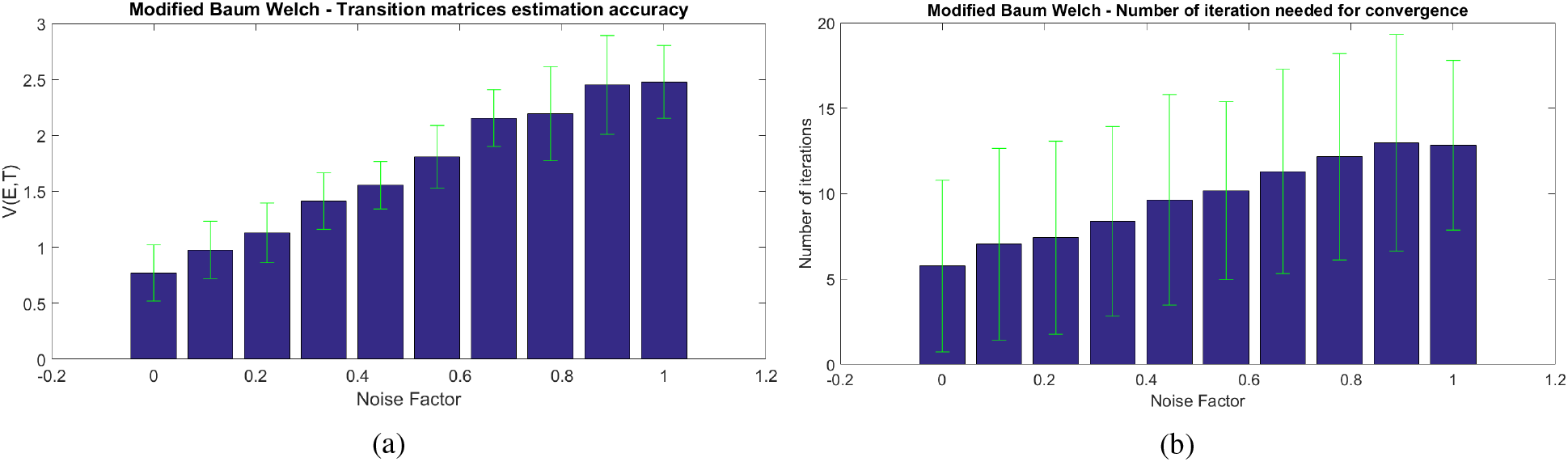
Top: The estimation parameter accuracy as a function of the value of noise to the perfect guess (Mean and STD). The metric used here is the variational distance. Bottom: the number of iterations needed for convergence as a function of the noise (Mean and STD).

There are several conditions in which using the BW algorithm on observations that were generated by the CA-HMM process may result in erroneous estimations due to the averaging over the different environments and, rewarded/unrewarded trials. For simplicity let us consider the case in which there are two policies and the transition and policy matrices are as follows:

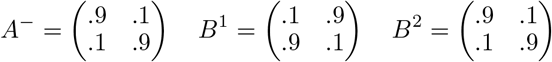

We generated a sequence of 1000 trials based on CA-HMM and estimated the model parameter using both BW and MBW, given that the guess of the transition matrices is [5., .5; .5,.5] for both BW and MBW. While the average score between the estimated emissions using MBW is 0.0851, the BW score distance is 1.68. The variational metric was used to calculate the score for both BW and MBW as follows:

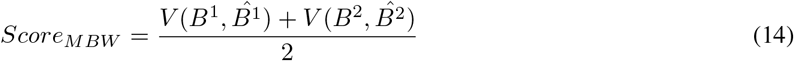

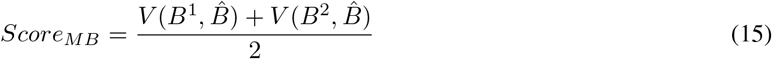

where 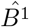 and 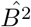 are the estimations returned by MBW and 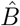 is the estimation returned by BW. The reported scores are the average over 50 trials (experiments) and using a tolerance factor of 10^*−*4^.

## 6 Related Work

Identifying the underlying policy that governs the observed behavior of an agent is an important task in many fields, including econometrics, neuroscience, game theory and machine-human interface.

The study of multi-agent systems (a sub-field that falls in the intersection between RL and game theory) is interested in reasoning about other agents’ intention and their decision-making process. This is usually a fundamental step toward selecting the optimal action in both cases of cooperation or competition. Another application is human and artificial agents behavioral modeling which is required for both successful human-AI collaboration and transparent (explainable) Artificial Intelligence (AI) [17, 18, 19, 20, 21].

A recent work by Unhelkar and Shah [17] considered the problem of inferring an agent’s decision-making model in a setup similar but not identical to the one presented in this paper. The decision-maker agent is assumed to be operating within a Markovian world with known dynamics (state transition function). Their approach first addresses the problem of identifying the presence of any latent state features given trajectories of observed behavior. Then, it estimates the policy of the agent given both the observed and latent features. However, there are several differences in the assumptions between their work than the current work: First, in this work, we assume that the set of features the agent considers is fully specified and known to the observer. Second, they assume that the agent employs a single stationary policy while in this work we assume that the agent may use a set of policies., thus the effort in this work is mainly directed in estimating the policies and their transition dynamics in the presence and absence of reward. Third, in [17] the observer has the ability to query the agent agent, while in this work, the observed behavioral trajectories is the only available information.

The *“Inverse reinforcement learning”* (IRL) problem described in [22] handles the case when the policy is fixed. IRL is defined as learning desirable behavior from a limited number of demonstrations. In general, the task of learning from an expert is called *apprenticeship learning* (also learning by watching, imitation learning, or learning from demonstration) [23, 24]. This setting is useful in applications where it may be difficult to write down an explicit reward function specifying exactly how different desiderata should be traded off, such as the task of driving. However, the policy estimation problem setup we are interested in is different from the IRL problem. First, in the IRL setting, the reward function is to be estimated, however, in the policy estimation problem, the reward function is fully known to the observer. Second, the expert policy in IRL is stationary, while the policy in the estimation problem can be dynamic.

In neuroscience, the interest in estimating the underlying dynamics of observed trial-by-trial data has gained increased attention in recent years [25], especially for modeling neural activity and behavior of animals with respect to input stimuli [26, 27, 1]. Here we will mention several traditionally used techniques for estimating underlying system dynamics by observing its output variables. The different approaches vary in their complexity, set of assumptions, and has different estimation challenges, therefore, the selection of the suitable approach is primarily governed by the aspect of the latent dynamics the researcher is mostly interested in.

Due to its enormous advantages, the scientific community has developed many extensions and generalizations of the basic HMM to handle the large variety of setups and assumptions on the underlying distribution of transitions and outputs that has emerged from the different applications. Reviewing the enormous body of work done on extending and adapting HMM is out of this paper scope, nevertheless, in the following, we will mention several related extensions. Infinite HMM has been proposed by in [28], non stationary HMM was developed in [16], Input-Output HMM was proposed by [29]. In many of these works developing the suitable parameter estimation method required adapting the Baum-Welch algorithm [16, 30] or in other cases developing a new identification approaches, however, most of these methods are based on the estimation-maximization (EM) algorithm or Monte Carlo based approaches, e.g. Monte Carlo Markov Chain [31].

The most popular models used to explain behavior are those derived from RL [8, 32, 1], which provides a method for an initially naive agent to achieve eventually optimal performance, by trial and error. As previously mentioned, the framework that describes this setup is referred to as MDP. Modeling the behavioral experiment as an MDP, one would need to estimate the policy of the behaving agent trial-by-trial by continuously observing the system state and the agent actions and assuming the underlying internal variables update procedure. In [33], the authors estimated the internal meta-parameters of the learning process, such as, the learning rate, the soft-max inverse temperature and discount factor of the assumed underlying Q-learning process using particle filter methods. They successfully applied the model for estimating the underlying processes of a learning monkey in a stochastic environment and show that the model could accurately estimate the animal behavior on a trial-by-trial basis.

A method for trial by trial behavior analysis of an animal in the plus maze, named Weighted attention model (WAM), was presented by Aluisi et al. in [34]. Unlike the previously reviewed approaches, and similar to our approach, this method is specifically interested in attentional set-shift. WAM learns by assigning a separate learning rule for the values of features of each dimension (e.g., for each color), reinforced after every experience. Decisions are made by comparing weighted averages of the learned values, factored by dimension specific weights. They showed that their approach captures mistakes made by the animal better than naive reinforcement estimation model.

## 7 Summary and Future Work

In this work, we aim at developing a method for identifying and measuring AST of an animal subject in the plus-maze, based on its observed behavior. Doing so required us to formulate the problem of estimating policy trajectory on a trial-by-trial basis while observing a behaving agent in a contextual bandit environment. The observed information is assumed to be generated by two dependent dynamical processes, one governs the environmental state transition and rewards, which dynamics are changing but known to the observer, and the other is a Markov process that controls the agent behavior centered around a latent variable representing the current agent strategy.

We show that while estimating the agent’s policy in the general case is difficult, imposing several sensible assumptions makes the solution computationally feasible and we relate it to the well known HMM framework. We propose an input depended and reward based HMM, we call “Context Aware HMM”. While imposing restriction on both the environment and the agent underlying policy dynamics, our approach allows identifying the different policies used by the agent and estimating the transition between those policies in a dynamic environment that allows non-stationary reward dynamics.

In future work, a deeper statistical analysis of the model should be performed to uncover important properties of the proposed framework, such as identifiability and stability, etc. In addition, we will explore a richer model which relaxes the stationary assumption to account for the changes in the transition probabilities during learning. The Baum-Welch algorithm has several drawbacks that our algorithm inherits, such as the sensitivity to the initial parameters as well as the convergence to a local minimum, therefore, in future work, we will examine alternative identification approaches for CA-HMM. It is advisable to assume that the transition probabilities between the different policies, in some cases, maintain some sort of relationship, hence, we could refine the CA-HMM model, to reflect this relationship and consequently reduce the number of model parameters. In addition, note that relaxing any of the model assumptions allows a rich avenue for future research.

## Appendix: Deriving the MBW from EM

In this section we derive the equations of the Modified Baum-Welch (MBW) algorithm presented in Section 4 from the general Expectation Maximization (EM) algorithm [35]. EM is an iterative method for finding maximum likelihood estimates of parameters in statistical models by alternating between performing an expectation and maximization steps until convergence. In the expectation step, the expected value of the log likelihood function is calculated with respect to the conditional distribution of the unobserved variable, given the observed variables under the current estimate *θ^q^*. In our case, following Equation 7:

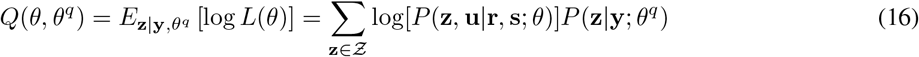

In the maximization step we find the parameter *θ* that maximize *Q*(*θ, θ^q^*):

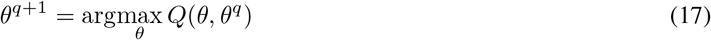

Therefore,

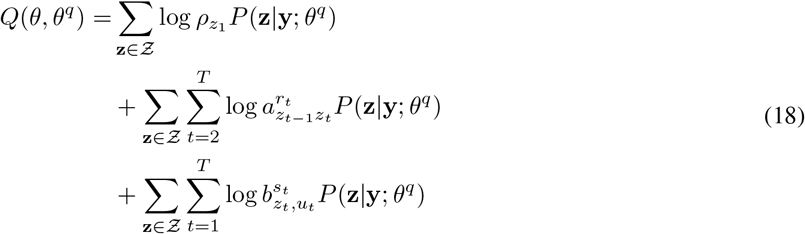

We will use the Langrangian multipliers technique to calculate the maximum in the domain restricting the transition and output probability to be a proper probability distribution (sums to 1). Let *𝓛*(*θ, θ^q^*) be the Lagrangian.

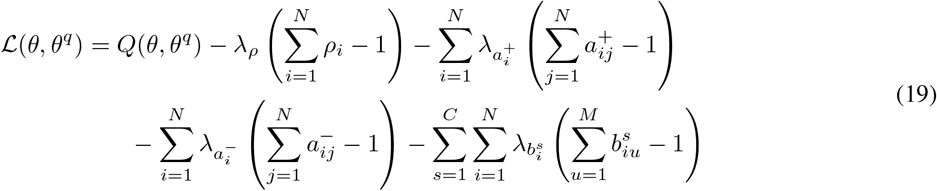

First lets focus our attention on the *ρ_i_*’s:

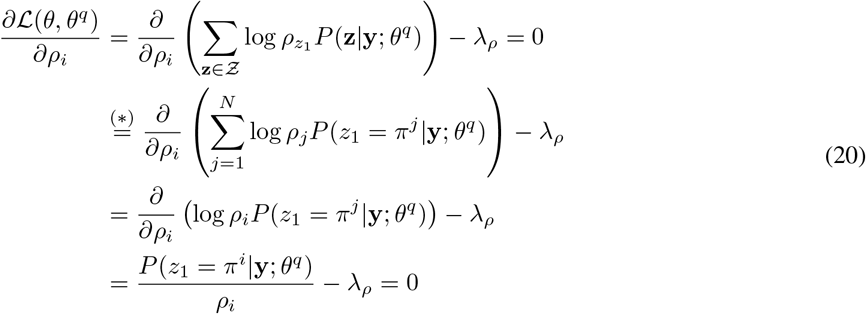

Transition (*) is a result of marginalizing out all *z*_*t≠1*_. The derivative with respect to the Lagrangian multiplier gives us the restriction as follows:

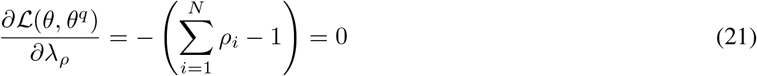

Now if we plug *ρ_i_* from equation 20 into Equation 21 we get:

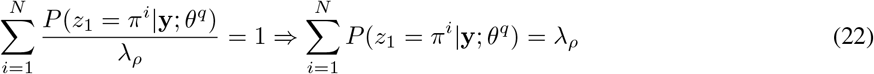

However, since 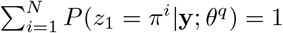 we get that *λ*_ρ_ = 1. Using Equation 20, again, for each parameter *ρ_i_*, the maximum point is obtained by:

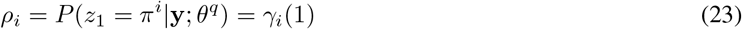

The last equality is by definition of *γ_i_*(*t*) in the previous section. This shows the derivation of Equation 12 for updating the initial state distribution.

We now follow a similar process for 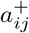 to derive Equation 10 in Section 4.

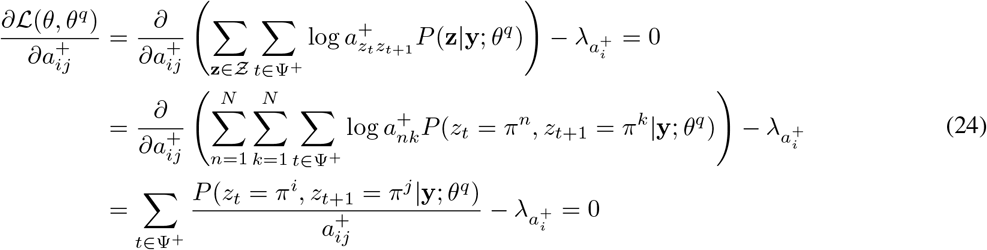

and the derivation with respect to the Lagrangian multipliers we get:

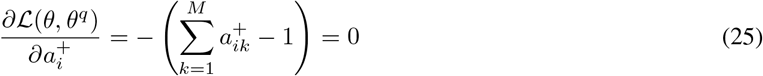

If we plug 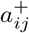 from equation 24 into Equation 25 we get:

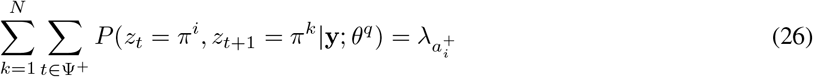

Plugging 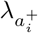 into Equation 24 we get:

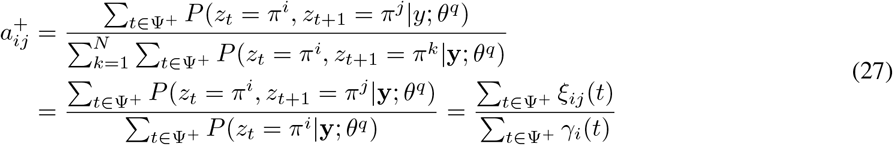

The last equality in Equation 27 is due to the definition of *ξ_ij_*(*t*) and *γ_i_*(*t*) An identical process can be easily carried out to derive 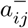.

Finally, we will show the same for 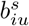. As before, let *I*(*x*) denote an indicator function.

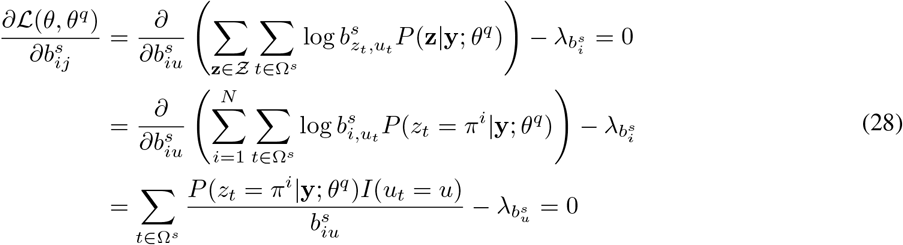

and

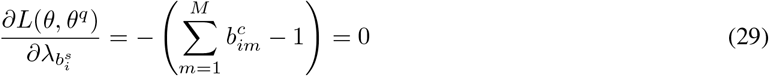

This now should come with no surprise:

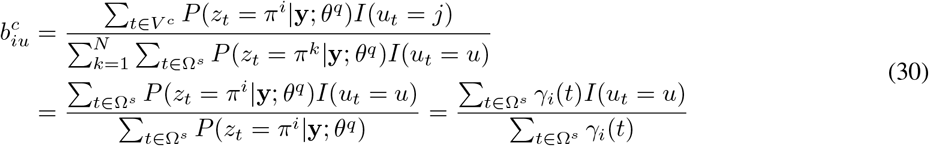

## References

[1] Greg Corrado and Kenji Doya. Understanding neural coding through the model-based analysis of decision making. Journal of Neuroscience, 27(31):8178–8180, 2007.

[2] Jillian M Heisler, Juan Morales, Jennifer J Donegan, Julianne D Jett, Laney Redus, and Jason C O’Connor. The attentional set shifting task: a measure of cognitive flexibility in mice. Journal of visualized experiments: JoVE, (96), 2015.

[3] Esta A Berg. A simple objective technique for measuring flexibility in thinking. The Journal of general psychology, 39(1):15–22, 1948.

[4] Alan H Nagahara, Tim Bernot, and Mark H Tuszynski. Age-related cognitive deficits in rhesus monkeys mirror human deficits on an automated test battery. Neurobiology of aging, 31(6):1020–1031, 2010.

[5] PL Rock, JP Roiser, WJ Riedel, and AD Blackwell. Cognitive impairment in depression: a systematic review and meta-analysis. Psychological medicine, 44(10):2029–2040, 2014.

[6] DT Stuss, B Levine, MP Alexander, J Hong, C Palumbo, L Hamer, KJ Murphy, and D Izukawa. Wisconsin card sorting test performance in patients with focal frontal and posterior brain damage: effects of lesion location and test structure on separable cognitive processes. Neuropsychologia, 38(4):388–402, 2000.

[7] Karl J Åström. Optimal control of markov processes with incomplete state information. Journal of Mathematical Analysis and Applications, 10(1):174–205, 1965.

[8] Richard S Sutton and Andrew G Barto. Reinforcement learning: An introduction, volume 1. MIT press Cambridge, 1998.

[9] John Langford and Tong Zhang. The epoch-greedy algorithm for multi-armed bandits with side information. In Advances in neural information processing systems, pages 817–824, 2008.

[10] Edward L Thorndike. The law of effect. The American Journal of Psychology, 39(1/4):212–222, 1927.

[11] Yingyao Hu and Matthew Shum. Nonparametric identification of dynamic models with unobserved state variables. Journal of Econometrics, 171(1):32–44, 2012.

[12] Leonard E Baum, Ted Petrie, George Soules, and Norman Weiss. A maximization technique occurring in the statistical analysis of probabilistic functions of markov chains. The annals of mathematical statistics, 41(1):164–171, 1970.

[13] Stephen E Levinson, Lawrence R Rabiner, and Man Mohan Sondhi. An introduction to the application of the theory of probabilistic functions of a markov process to automatic speech recognition. Bell System Technical Journal, 62(4):1035–1074, 1983.

[14] Frederick Jelinek, Lalit Bahl, and Robert Mercer. Design of a linguistic statistical decoder for the recognition of continuous speech. IEEE Transactions on Information Theory, 21(3):250–256, 1975.

[15] Solomon Kullback and Richard A Leibler. On information and sufficiency. The annals of mathematical statistics, 22(1):79–86, 1951.

[16] Bongkee Sin and Jin H Kim. Nonstationary hidden markov model. Signal Processing, 46(1):31–46, 1995.

[17] Vaibhav V Unhelkar and Julie A Shah. Learning models of sequential decision-making without complete state specification using bayesian nonparametric inference and active querying. 2018.

[18] Vincent Aleven, Jonathan Sewall, Octav Popescu, Franceska Xhakaj, Dhruv Chand, Ryan Baker, Yuan Wang, George Siemens, Carolyn Rosé, and Dragan Gasevic. The beginning of a beautiful friendship? intelligent tutoring systems and moocs. In International Conference on Artificial Intelligence in Education, pages 525–528. Springer, 2015.

[19] Aditi Ramachandran, Chien-Ming Huang, and Brian Scassellati. Give me a break!: Personalized timing strategies to promote learning in robot-child tutoring. In Proceedings of the 2017 ACM/IEEE International Conference on Human-Robot Interaction, pages 146–155. ACM, 2017.

[20] Andrea Thomaz, Guy Hoffman, Maya Cakmak, et al. Computational human-robot interaction. Foundations and Trends® in Robotics, 4(2-3):105–223, 2016.

[21] Bradley Hayes and Julie A Shah. Improving robot controller transparency through autonomous policy explanation. In Proceedings of the 2017 ACM/IEEE international conference on human-robot interaction, pages 303–312. ACM, 2017.

[22] Pieter Abbeel and Andrew Y Ng. Inverse reinforcement learning. In Encyclopedia of machine learning, pages 554–558. Springer, 2011.

[23] Pieter Abbeel and Andrew Y Ng. Apprenticeship learning via inverse reinforcement learning. In Proceedings of the twenty-first international conference on Machine learning, page 1. ACM, 2004.

[24] Brenna D Argall, Sonia Chernova, Manuela Veloso, and Brett Browning. A survey of robot learning from demonstration. Robotics and autonomous systems, 57(5):469–483, 2009.

[25] Nathaniel D Daw. Trial-by-trial data analysis using computational models. Decision making, affect, and learning: Attention and performance XXIII, 23:3–38, 2011.

[26] Anne C Smith and Emery N Brown. Estimating a state-space model from point process observations. Neural computation, 15(5):965–991, 2003.

[27] Yingzhuo Zhang, Noa Malem-Shinitski, Stephen A Allsop, Kay Tye, and Demba Ba. Estimating a separably-markov random field (smurf) from binary observations. arXiv preprint arXiv:1709.09723, 2017.

[28] Matthew J Beal, Zoubin Ghahramani, and Carl E Rasmussen. The infinite hidden markov model. In Advances in neural information processing systems, pages 577–584, 2002.

[29] Yoshua Bengio and Paolo Frasconi. An input output hmm architecture. In Advances in neural information processing systems, pages 427–434, 1995.

[30] Paul M Baggenstoss. A modified baum-welch algorithm for hidden markov models with multiple observation spaces. In Acoustics, Speech, and Signal Processing, 2000. ICASSP’00. Proceedings. 2000 IEEE International Conference on, volume 2, pages II717–II720. IEEE, 2000.

[31] Michael David Escobar. Estimating the means of several normal populations by nonparametric estimation of the distribution of the means. 1990.

[32] Kenji Doya. Reinforcement learning: Computational theory and biological mechanisms. HFSP journal, 1(1):30, 2007.

[33] Kazuyuki Samejima, Kenji Doya, Yasumasa Ueda, and Minoru Kimura. Estimating internal variables and paramters of a learning agent by a particle filter. In NIPS, pages 1335–1342, 2003.

[34] Flavia Aluisi, Anna Rubinchik, and Genela Morris. Animal learning in a multidimensional discrimination task as explained by dimension-specific allocation of attention. Frontiers in Neuroscience, 12, 2018.

[35] Arthur P Dempster, Nan M Laird, and Donald B Rubin. Maximum likelihood from incomplete data via the em algorithm. Journal of the royal statistical society. Series B (methodological), pages 1–38, 1977.

